# Genome-wide association and functional studies identify 46 novel loci for alcohol consumption and suggest common genetic mechanisms with neuropsychiatric disorders

**DOI:** 10.1101/453332

**Authors:** Evangelos Evangelou, He Gao, Congying Chu, Georgios Ntritsos, Paul Blakeley, Andrew R. Butts, Raha Pazoki, Hideaki Suzuki, Fotios Koskeridis, Andrianos M. Yiorkas, Ibrahim Karaman, Joshua Elliott, Stefanie Aeschbacher, Traci M. Bartz, Sebastian E. Baumeister, Peter S. Braund, Michael R. Brown, Jennifer A. Brody, Toni-Kim Clarke, Niki Dimou, Jessica D. Faul, Georg Homuth, Anne U. Jackson, Katherine A. Kentistou, Peter K. Joshi, Rozenn N. Lemaitre, Penelope A. Lind, Leo-Pekka Lyytikäinen, Massimo Mangino, Yuri Milaneschi, Christopher P. Nelson, Ilja M. Nolte, Mia-Maria Perälä, Ozren Polasek, David Porteous, Scott M. Ratliff, Jennifer A. Smith, Alena Stančáková, Alexander Teumer, Samuli Tuominen, Sébastien Thériault, Jagadish Vangipurapu, John B. Whitfield, Alexis Wood, Jie Yao, Bing Yu, Wei Zhao, Dan E. Arking, Juha Auvinen, Chunyu Liu, Minna Männikkö, Lorenz Risch, Jerome I. Rotter, Harold Snieder, Juha Veijola, Alexandra I. Blakemore, Michael Boehnke, Harry Campbell, David Conen, Johan G. Eriksson, Hans J. Grabe, Xiuqing Guo, Pim van der Harst, Catharina A. Hartman, Caroline Hayward, Andrew C. Heath, Marjo-Riitta Jarvelin, Mika Kähönen, Sharon LR Kardia, Michael Kühne, Johanna Kuusisto, Markku Laakso, Jari Lahti, Terho Lehtimäki, Andrew M. McIntosh, Karen L. Mohlke, Alanna C. Morrison, Nicholas G. Martin, Albertine J. Oldehinkel, Brenda WJH Penninx, Bruce M. Psaty, Olli T. Raitakari, Igor Rudan, Nilesh J. Samani, Laura J. Scott, Tim D. Spector, Niek Verweij, David R. Weir, James F. Wilson, Daniel Levy, Ioanna Tzoulaki, Jimmy D. Bell, Paul Matthews, Adrian Rothenfluh, Sylvane Desrivières, Gunter Schumann, Paul Elliott

**Author notes:** Equal contribution (Evangelos Evangelou, He Gao, Congying Chu, Georgios Ntritsos, Jimmy D. Bell, Paul Matthews, Adrian Rothenfluh, Sylvane Desrivières, Gunter Schumann, Paul Elliott). Corresponding authors: Gunter Schumann and Paul Elliott.

## Abstract

Excessive alcohol consumption is one of the main causes of death and disability worldwide. Alcohol consumption is a heritable complex trait. We conducted a genome-wide association study (GWAS) of alcohol use in ~480,000 people of European descent to decipher the genetic architecture of alcohol intake. We identified 46 novel, common loci, and investigated their potential functional significance using magnetic resonance imaging data, gene expression and behavioral studies in *Drosophila*. Our results identify new genetic pathways associated with alcohol consumption and suggest common genetic mechanisms with several neuropsychiatric disorders including schizophrenia.

---

Excessive alcohol consumption is a major public health problem that is responsible for 2.2% and 6.8% age-standardized deaths for women and men respectively^1^. Most genetic studies of alcohol use focus on alcohol dependency, although the burden of alcohol-related disease mainly reflects a broader range of alcohol consumption behaviors in a population^2^. Small reductions in alcohol intake could have major public health benefits; a recent study reported that even moderate daily alcohol may have significant impact on mortality^3^.

Alcohol consumption is a heritable complex trait^4^, but genetic studies to date have identified only a small number of robustly associated genetic variants ^5-8^. These include variants in the aldehyde dehydrogenase gene family, a group of enzymes that catalyze the oxidation of aldehydes^9^, including a cluster of genes on chromosome 4q23 (*ADH1B, ADH1C, ADH5, ADH6, ADH7*)^6^.

Here, we report a GWAS meta-analysis of alcohol intake (g/day) among people of European ancestry drawn from UK Biobank (UKB)^10^, the Alcohol Genome-Wide Consortium (AlcGen) and the Cohorts for Heart and Aging Research in Genomic Epidemiology Plus (CHARGE+) consortia. Briefly, UKB is a prospective cohort study of ~500,000 individuals recruited between the ages of 40-69 years. Participants were asked to report their average weekly and monthly alcohol consumption through a self-completed touchscreen questionnaire^10^. Based on these reports, we calculated the gram/day (g/d) alcohol intake (**Online Methods**). Participants were genotyped using a customized array with imputation from the Haplotype Reference Consortium (HRC) panel^11^, yielding ~7 million common single nucleotide polymorphisms (SNPs) with minor allele frequency (MAF) ≥ 1% and imputation quality score [INFO] ≥ 0.1. After quality control (QC) and exclusions (**Online Methods**) we performed GWAS of alcohol consumption using data from 404,731 UKB participants of European descent under an additive genetic model (**Online Methods and Supplementary Table 1**). We found that genomic inflation in the UKB analysis was λ_GC_=1.45, but did not adjust for inflation as the LD score regression intercept was 1.05, indicating that this was due to polygenicity rather than to population stratification^12^. The estimated SNP-wide heritability of alcohol consumption in the UKB data was 0.09.

We also carried out GWAS in 25 independent studies from the AlcGen and CHARGE+ consortia including 76,111 participants of European descent for which alcohol g/d could be calculated (**Supplementary Table 2**). Various arrays were used for genotyping, with imputations performed using either the 1,000 Genomes Reference Panel or the HRC platforms (**Supplementary Table 3**). After QC, we applied genomic control at the individual study level and obtained summary results for ~7 million SNPs with imputation quality score ≥ 0.3 (**Online Methods**).

We combined the UKB, AlcGen and CHARGE+ results using a fixed effects inverse variance weighted approach for a total of 480,842 individuals^13^. To maximize power, we performed a single-stage analysis to test common SNPs with MAF ≥ 1%. We set a stringent *P*-value threshold of *P* < 5 × 10^-9^ to denote significance in the combined meta-analysis^14^, and required signals to be significant at *P* < 5 × 10^-7^ in UKB, with same direction of effect in UKB and AlcGen plus CHARGE+, to minimize false positive findings. We excluded SNPs within 500kb of variants reported as genome-wide significant in previous GWAS of alcohol consumption^5,6^, identified novel loci by requiring SNPs to be independent of each other (LD r^2^ < 0.1), and selected the sentinel SNP within each locus according to lowest *P*-value (**Online Methods**).

We then tested for correlations of alcohol-associated SNPs with Magnetic Resonance Imaging (MRI) phenotypes of brain, heart and liver, and gene expression. Associations of the sentinel SNPs with other traits/diseases were investigated and *Drosophila* mutant models used to test for functional effects on ethanol-induced behavior.

## Results

Our meta-analysis identified 46 novel loci associated with alcohol consumption (log transformed g/day) (**Fig. 1 and Table 1**). We discovered eight additional variants in the combined analysis at nominal genome-wide significance (*P* < 1 × 10^-8^) that may also be associated with alcohol intake (**Supplementary Table 4**). The most significantly associated variant, rs1991556 (*P* < 4.5 × 10^-23^), is an intronic variant in *MAPT* gene that encodes the microtubule-associated protein tau, and was found through Phenoscanner not only to be associated with dementia^15^ and Parkinson’s disease^16,17^, but also with neuroticism, schizophrenia^18^ and other conditions^19-21^ (**Online Methods, Fig. 2 and Supplementary Table 5**). The second most significantly associated variant is rs1004787 (*P* < 6.7 × 10^-17^), near *SIX3* gene, which encodes a member of the sine oculis homeobox transcription factor family involved in eye development^22^. The third SNP is rs13107325 (*P* < 1.3 × 10^-15^), a missense SNP in *SLC39A8*, a gene that encodes a member of the SLC39 family of metal ion transporters, which has been associated with schizophrenia^23^ as well as inflammatory bowel disease, cardiovascular and metabolic phenotypes ^24^ ^25-27^ in previous GWAS (**Fig. 2 and Supplementary Table 5**).

**Figure 1.**
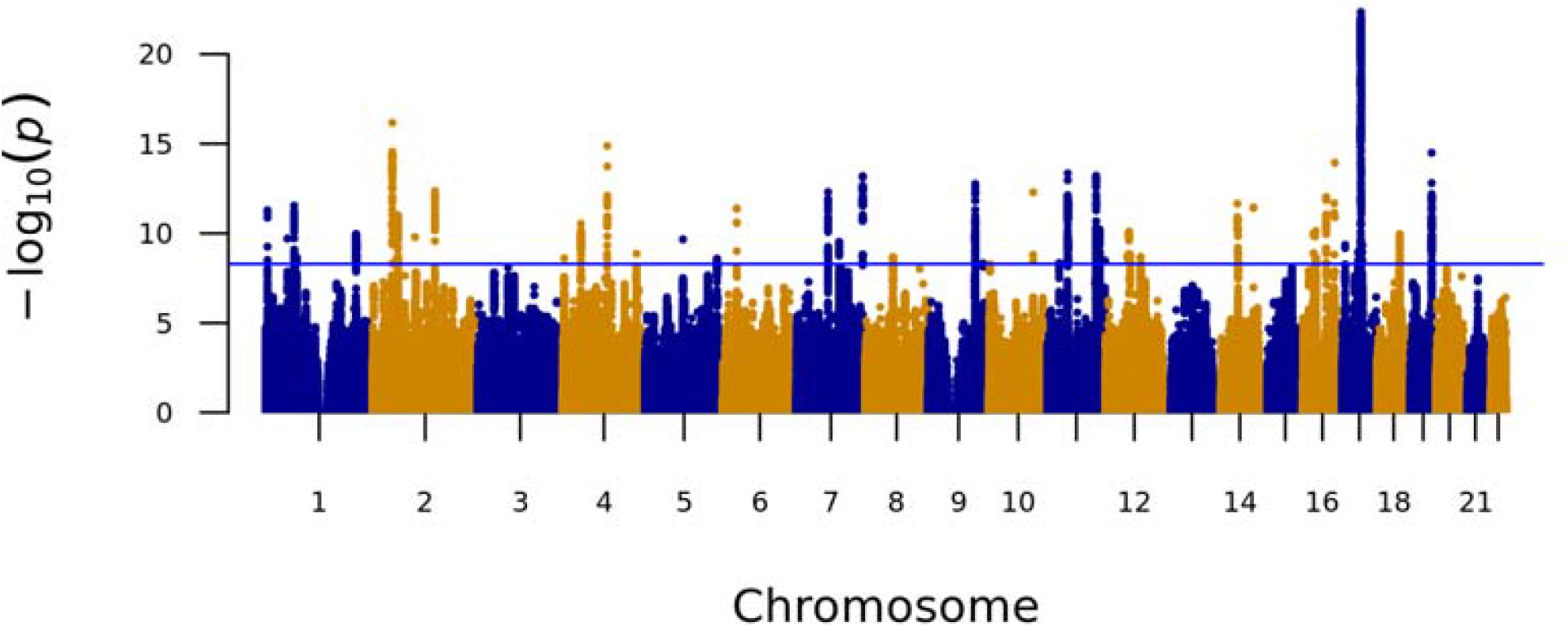
Manhattan plot showing *P*-values from discovery genome-wide association meta-analysis with alcohol intake (g/d) among 480,842 individuals across UK Biobank, AlcGen and CHARGE+, excluding known variants. The *P*-value was computed using inverse variance fixed effects models. The y axis shows the –log_10_ P values and the × axis shows their chromosomal positions. Horizontal blue line represents the threshold of *P* = 5 × 10^-9^.

**Figure 2.**
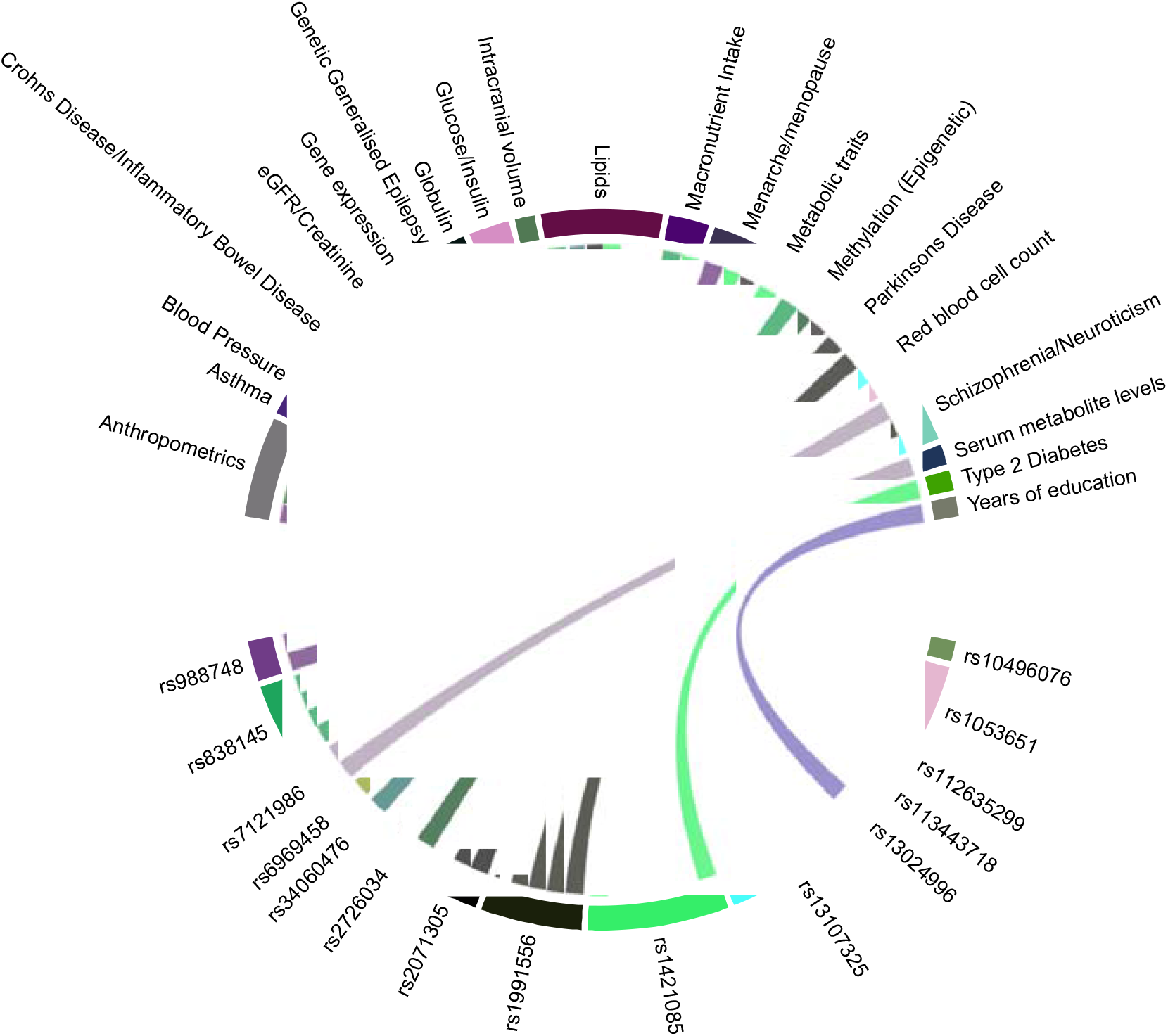
Association of alcohol intake loci with other traits. Plot shows results from associations with other traits which were extracted from the PhenoScanner database for the 46 novel sentinel SNPs including proxies in Linkage Disequilibrium (r^2^ ≥ 0.8) with genome-wide significant associations.

**Table 1.**
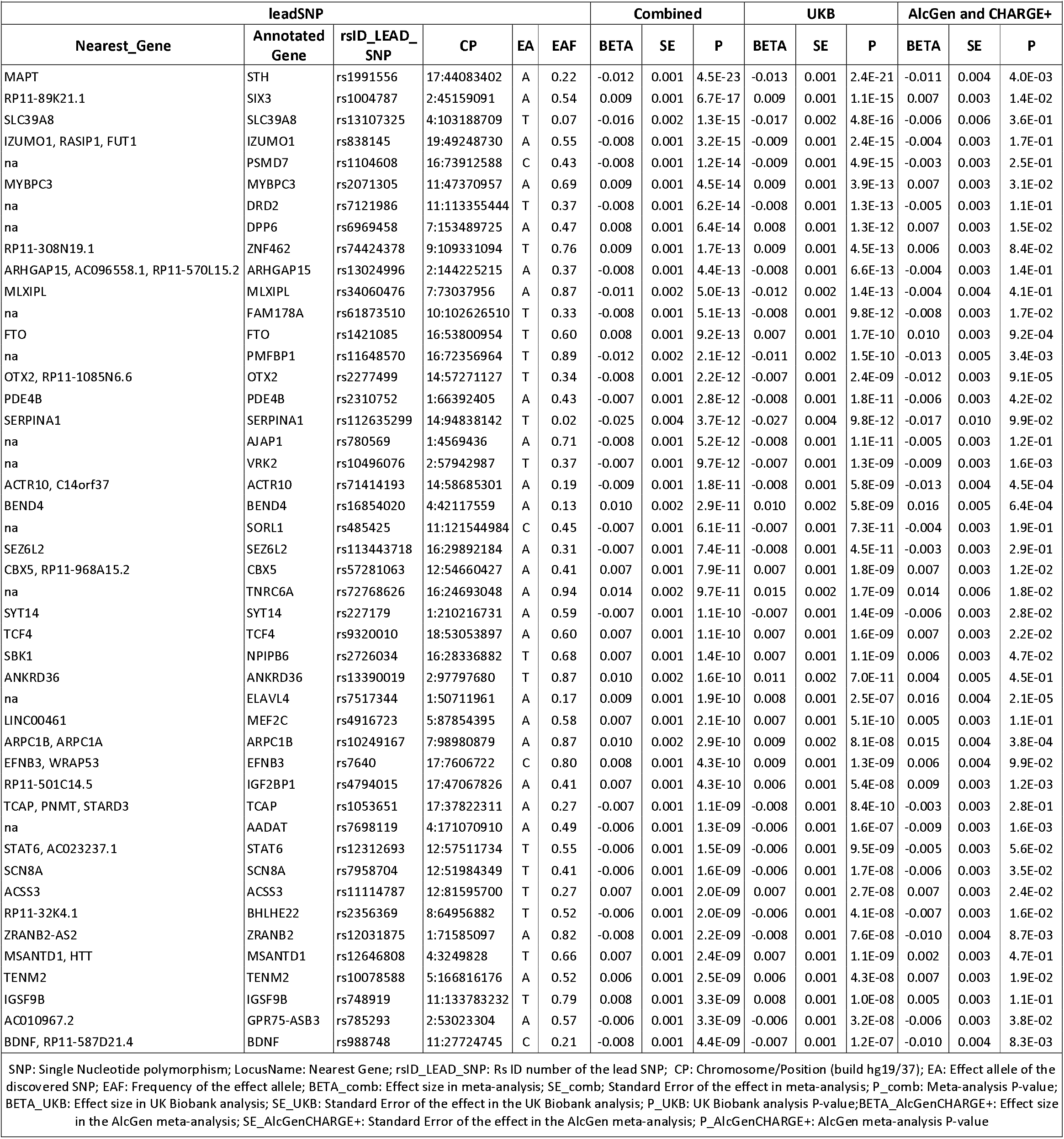
Association results of 46 novel alcohol variants identified through the meta-analysis of UK Biobank and AlcGen and CHARGE+. Results are ordered by *P*-value of combined analysis.

Another of our most significant variants, an intronic SNP rs7121986 (*P* < 6.2 × 10^-14^) in *DRD2*, encodes the dopamine receptor D2 that has been associated with cocaine addiction, neuroticism and schizophrenia^18^. We also found significant associations with SNP rs988748 (*P* < 4.4 × 10^-9^) in the gene encoding BDNF (brain-derived neurotrophic factor) and rs7517344, which is near *ELAVL4* (*P* = 2.0 × 10^-10^), the gene product of which is involved in *BDNF* regulation^28^. Previous studies have suggested that variation in *BDNF* is a genetic determinant of alcohol consumption and that alcohol consumption modulates *BDNF* expression^29^.

Additionally, we found association of alcohol consumption with SNP rs838145 (*P* < 3.2 × 10^-15^), which has been associated with macronutrient intake in a previous GWAS^30^. This variant is localized to *IZUMO1*, a locus of around 50kb that spans a number of genes including *FGF21*, whose gene product *FGF21* is a liver hormone involved in the regulation of alcohol preference, glucose and lipid metabolism^31^. We previously reported significant association of alcohol intake with SNP rs11940694 in *KLB*, an obligate receptor of *FGF21* in the brain^5^, and strongly replicated that finding here (*P* = 3.3 × 10^-68^).

As well as variants in *KLB*, we found support (*P* < 1 × 10^-5^) for association of common variants in all four of the other previously reported alcohol intake-related loci (**Supplementary Table 6**). These replicated loci include SNP rs6943555 in *AUTS2* (*P* = 2.9 × 10^-6^) and variants in the alcohol dehydrogenase locus (lowest *P* = 1.2 × 10^-125^). In addition, we found a novel alcohol intake-related SNP rs1421085 in *FTO* in high LD (r^2^ = 0.92) with a variant reported previously as genome-wide significant for association with alcohol dependence^32^.

Conditional analysis using Genome-wide Complex Trait Analysis (GCTA) did not reveal any independent secondary signals related to alcohol consumption. Among ~14,000 individuals in the independent Airwave cohort^33^ (**Online Methods**), 7% of the variance in alcohol consumption was explained by the novel and known common variants. Using weights from our analysis, we constructed an unbiased weighted genetic risk score (GRS) in Airwave (**Online Methods**) and found a strong association of the novel and known variants on alcohol consumption levels (*P* = 2.75 × 10^-14^), with mean difference in sex-adjusted alcohol intake of 2.6 g/d comparing the top vs the bottom quintile of the GRS (**Supplementary Table 7**).

### Associations with MRI imaging phenotypes

We performed single-SNP analyses of the imaging phenotypes in UKB (**Online Methods**) to investigate associations of our novel variants with MRI of brain (N=9,702), heart (N=10,706) and liver (N=8,479). With Bonferroni correction (corrected *P*-value 6.6 × 10^-6^, corresponding to 0.05/46 SNPs*164 imaging phenotypes), we found significant positive associations between rs13107325 and the volumes of multiple brain regions; the strongest associations were with putamen (left: *P* = 2.5 × 10^-45^, right: *P* = 2.8 × 10^-47^), ventral striatum (left: *P* = 9.5 × 10^-53^, right: *P* = 9.6 × 10^-51^) and cerebellum (strongest association for left I-IV volume; *P* = 1.2 × 10^-9^) (**Supplementary Table 8**); similar findings were also recently reported in a GWAS on brain imaging in UKB34. The other significant association was for rs1991556 with the parahippocampal gyrus (*P* = 1.2 × 10^-6^).

We then tested these brain regions for association with alcohol consumption and found a significant effect for the left (*P* = 2.0 × 10^-4^) and right (*P* = 2.6 × 10^-4^) putamen. Finally, we used data from N= 8,610 individuals and performed a mediation analysis using a standard three-variable path model, bootstrapping 10,000 times to calculate the significance of the mediation effect of putamen volume for genetic influences on alcohol consumption (**Online Methods**). We found evidence that the effect of SNP rs13107325 in *SLC39A8* on alcohol intake is partially mediated via its association with left (beta=-0.27; *P* = 1.9 × 10^-3^) and right (beta=-0.26; *P* = 1.7 × 10^-3^) putamen volume (**Fig. 3 and Supplementary Table 9**).

**Figure 3.**
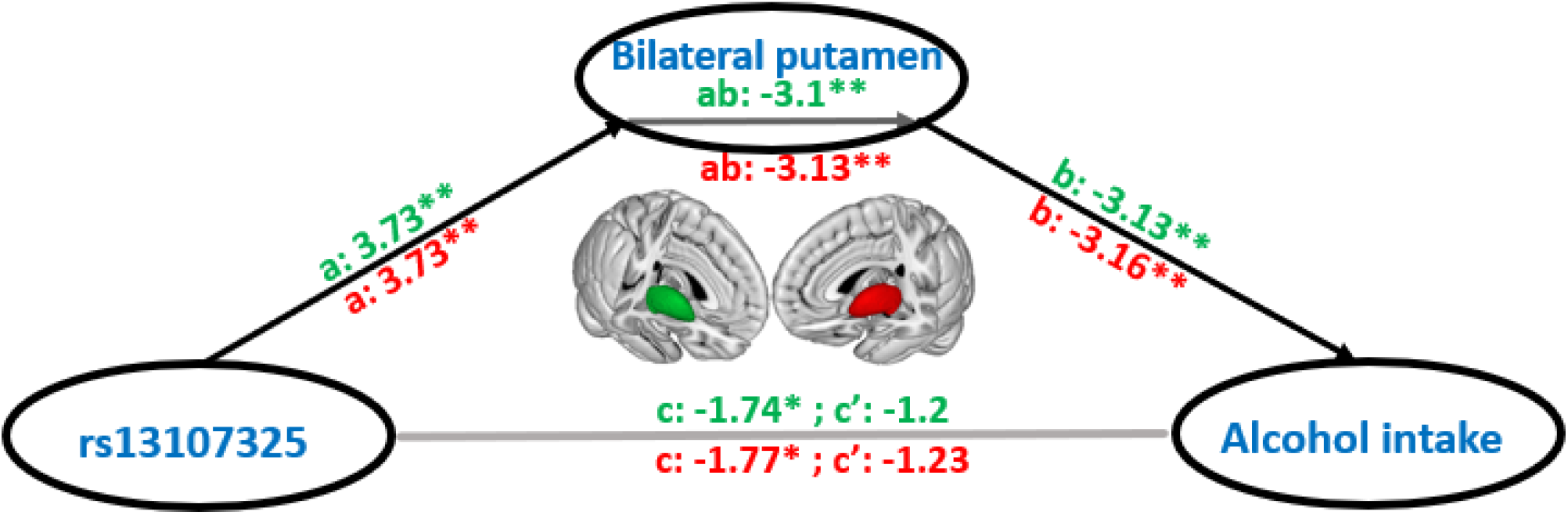
Mediation effect of bilateral putamen on the relationship between SNP rs13107325 and alcohol intake. Left putamen is indicated by the green color whereas the right putamen by the red. a presents the association between rs13107325 and putamen, b is the association between putamen and alcohol consumption, c the association of rs13107325 and alcohol consumption, c’ is the association between rs13107325 and alcohol consumption after excluding the effect of putamen, and ab is the mediation effect. The significance of the effect is based on bootstrapping. We provide the z-statistic for each relationship combined with *P*-values (** as *P* < 0.005, * as *P* < 0.1).

We did not find any significant associations of novel SNPs with either cardiac (left ventricular mass or end diastolic volume or right ventricular end diastolic volume) (**Supplementary Table 10**) or liver fat measures on MRI (**Supplementary Table 11**), after adjustment for multiple testing.

### Effects of SNPs on gene expression

We carried out expression quantitative trait loci eQTL analyses using the Genotype-Tissue Expression (GTEx) and the UK Brain Expression Consortium (UKBEC) datasets; 34 of the 53 novel and known SNPs associated with alcohol consumption have a significant effect on gene expression in at least one tissue, including 33 SNPs that affected gene expression in the brain (**Supplementary Tables 12 and 13, and Supplementary Fig. 1-4**). We found that the most significant eQTLs often do not involve the nearest gene and that several of the SNPs affect expression of different genes in different tissues (**Supplementary Fig. 4**). For example, SNP rs1991556 in the *MAPT* gene affects expression of 33 genes overall, with most significant effects on the expression of the non-protein coding genes *CRHR1-IT1* (also known as *C17orf69* or *LINC02210*) and *LRRC37A4P*, near *MAPT*, across a wide range of tissues including brain, adipose tissue and skin (*P* = 7.2 × 10^-126^ to *P* = 2.5 × 10^-6^) (**Supplementary Fig. 4**). Similarly, the A-allele at SNP rs2071305 within *MYBPC3* affects the expression of several genes and is most significantly associated with increased expression of *C1QTNF4* across several tissues (*P* = 1.9 × 10^-25^ to *P* = 8.4 × 10^-5^).

Several of these eQTLs were found to affect expression of genes known to be involved in reward and addiction. SNP rs1053651 in the *TCAP-PNMT-STARD3* gene cluster affects expression of the *PPP1R1B* gene (also known as *DARPP-32*) which encodes a protein that mediates the effects of dopamine in the mesolimbic reward pathway^35^. Other known addiction-related genes include *ANKK1* and *DRD2* (affected by SNP rs7121986) implicated in alcohol and nicotine dependence^36,37^, CRHR1 (affected by SNP rs1991556) involved in stress-mediated alcohol dependence^38,39^ and *PPM1G* (SNP rs1260326) whose epigenetic modification was reported to be associated with alcohol abuse^40^.

Over-representation enrichment analyses based on functional annotations and disease-related terms indicated that genes whose expressions are affected by the identified eQTLs are most significantly enriched for terms related to abdominal cancers (n = 91), motor function (n= 5) and cellular homeostasis (n= 22) (**Supplementary Fig 5**).

### Other traits and diseases

Using LD score regression^12^, we assessed genetic correlations between alcohol consumption and 235 complex traits and diseases from publicly available summary GWAS statistics (**Online Methods and Supplementary Table 14**). The strongest positive genetic correlations based on false discovery rate *P* < 0.02 were found for smoking (r_g_ = 0.42, *P* = 1.0 × 10^-23^) and HDL cholesterol levels (r_g_ = 0.26, *P* = 5.1 × 10^-13^). We also found negative correlations for sleep duration (r_g_ = −0.14, *P* = 3.8 × 10^-7^) and fasting insulin levels (r_g_ = −0.25, *P* = 4.5 × 10^-6^). A significant genetic correlation was also found with schizophrenia (r_g_ = 0.07, *P* = 3.9 × 10^-3^) and bipolar disorder (r_g_ = 0.15, *P* = 5.0 × 10^-4^) (**Supplementary Table 14**). Over-representation enrichment analysis using WebGestalt^41^ showed that our list of novel and known variants are significantly enriched in several diseases and traits including developmental disorder in children (*P* < 7.3 × 10^-5^), epilepsy (*P* < 1.4 × 10^-4^), heroin dependence (*P* = 5.7 × 10^-4^) and schizophrenia (*P* < 8.4 × 10^-4^) (**Supplementary Fig. 6**). The result of Mendelian randomization analysis (**Online methods**) to assess a potential causal effect of alcohol on schizophrenia risk, using the inverse variance weighted approach, was not significant (*P* = 0.089), with large heterogeneity of the estimates of the tested variants.

### Functional studies in *Drosophila*

Based on our GWAS and brain imaging findings we took forward SNP rs13107325 in *SLC39A8* (alias *Zip8* gene) for additional testing in *Drosophila*, which employ conserved mechanisms to modulate ethanol-induced behaviors^42,43^. First, we overexpressed human *Zip8* using a Gal4-driver that included expression in neurons involved in multiple ethanol-induced behaviors^43^. Flies carrying icsGal4/+ *UAS-hZip8*/+ showed a slight, but significant, resistance to ethanol-induced sedation compared to control flies (*P* = 0.026; N = 16 per genotype). Ethanol tolerance, induced with repeat exposures spaced by a 4-hour recovery, was unchanged in these flies (Fig. 4a). We next used the same Gal4-driver to knock down the endogenous *Drosophila* ortholog of *hZip8*, namely *dZip71B*. This caused the flies to display naïve sensitivity to ethanol-induced sedation, and in addition, these flies developed greater tolerance to ethanol upon repeat exposure (*P* = 0.0003; N = 8 per genotype) (Fig. 4b). To corroborate this phenotype, we then tested flies transheterozygous for two independent transposon-insertions in the middle of the *dZip71B* gene (**Supplementary Fig. 7**) and found that these *dZip71B^Mi / MB^* flies also displayed naïve sensitivity (*P* = 0.006) and increased ethanol-induced tolerance (*P* = 0.032), compared to controls (N = 8 each) (Fig. 4c).

**Figure 4.**
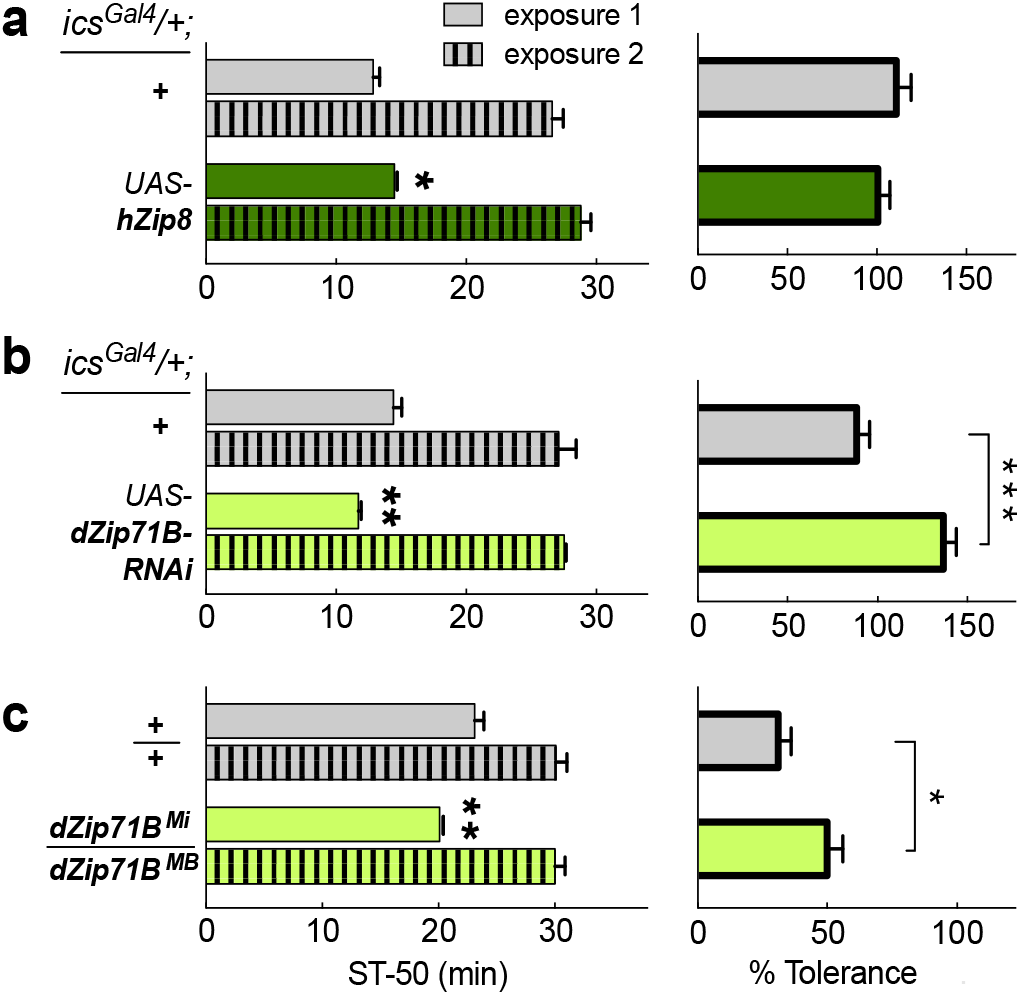
Comparison of *Zip8* alcohol phenotypes in *Drosophila*. Flies were exposed to 100/50 Ethanol/Air vapor for 30 min for exposure 1, and the time to 50% loss of righting was determined (ST-50, sedation time). After recovery on food for 4 hours, flies were re-exposed to the same vapors, and the second ST-50 recorded (left side). The resulting increase in ST-50, i.e. tolerance, is shown on the right. In a) overexpressed human *hZIP8* in ics-expressing cells flies are compared against controls whereas in b) knockdown of the fly ortholog *dZip71B* is compared against controls. In c) flies carrying two transposon insertions in the endogenous *dZip71B* gene are compared against controls. Significance levels: ***P <0.001, **P <0.01, **P <0.05. Actual *P*-values are presented in the text

## Discussion

Our discovery utilizing data on common variants from over 480,000 people of European descent has greatly extended our knowledge of the genetic architecture of alcohol intake, increasing the number of loci by nearly 10-fold to 46. We found loci involved in neuropsychiatric conditions such as schizophrenia, Parkinson’s disease and dementia, as well as *BDNF* where gene expression is affected by alcohol abuse. Our findings illustrate that large-scale studies of genetic associations with alcohol intake in the general population, rather than in alcohol dependency alone, can provide new insights into genetic mechanisms regulating alcohol consumption.

We highlight the role of the highly pleiotropic *MAPT* and *SLC39A8* genes in the genetics of alcohol consumption. *MAPT* plays a key role in tau-associated dementia^44^ and both genes are also implicated in other neuropsychiatric conditions including neuroticism, schizophrenia and Parkinson’s disease^16-18^. The *SLC39A8* gene encodes a member of the SLC39 family of metal ion transporters. The encoded protein is glycosylated and found in plasma membrane and mitochondria, and is involved in the cellular transport of zinc, modulation of which could affect microglial inflammatory responses^45^. Our gain- and loss-of-function studies in *Drosophila* indicate a potential causal role of *SLC39A8* in alcohol drinking behavior. The MRI brain imaging demonstrates a significant association of SNP rs13107325 in the *SLC39A8* gene and putamen volume differences, and these structural differences appear to partially mediate associations of rs13107325 with alcohol consumption. The putamen has been associated with alcohol consumption and the withdrawal syndrome after chronic administration to rodents and non-human primates^46^. Putamen volume differences have also been associated with both schizophrenia and psychosis^47,48^ and robust association between SNP rs13107325 in *SLC39A8* and schizophrenia was reported in a previous GWAS^23^.

We also report SNP rs7121986 near *DRD2* as a novel alcohol intake variant in GWAS. The gene product of *DRD2*, D2 dopamine receptor, is a G protein-coupled receptor on post-synaptic dopaminergic neurons that has long been implicated in alcoholism^49^. In addition, we identify SNP rs988748 in *BDNF* as a novel alcohol intake variant; BDNF expression is differentially affected by alcohol exposure in animal models^50,51^. Both genes (along with *PPP1R1P*) are centrally involved in reward-mediating mesocortico-limbic pathways and both are implicated in the development of schizophrenia. For example, there is a robust GWAS association between schizophrenia and SNP rs4938021 in *DRD2* (in perfect LD with our novel alcohol intake-related variant rs7121986) and *DRD2* appears to be pivotal in network analyses of genes involved in schizophrenia^52^. Taken together, our results suggest that there are shared genetic mechanisms between the regulation of alcohol intake and susceptibility to schizophrenia, as well as other neuropsychiatric disorders. In this regard, large prospective epidemiological studies report a three-fold risk of schizophrenia in relation to alcohol abuse^53^.

We previously reported genome-wide significant associations of alcohol intake with *KLB*, and identified a liver-brain axis linking the liver hormone *FGF21* with central regulation of alcohol intake involving β-Klotho receptor (the gene product of *KLB*) in the brain^5^. Here, we identify a significant variant near *FGF21* gene and strongly replicate the previously reported *KLB* gene variant, strengthening the genetic evidence for the importance of this pathway in regulating alcohol consumption.

The LD score regression analysis showed a positive genetic correlation between alcohol consumption, smoking and HDL cholesterol levels. This confirms previous findings that reported an almost identical genetic correlation of alcohol consumption with number of cigarettes per day^54^. Furthermore, the observed genetic correlation with HDL levels is consistent with previous observations of an association between alcohol consumption and HDL55,56, including results of a Mendelian randomization study that suggested a possible causal role linking alcohol intake with increased HDL levels^57^. Finally, we found a genetic correlation (inverse) between sleep duration and alcohol consumption, an association previously reported only in a few small epidemiological studies^58^. We could not test for a genetic association between alcohol and risk of alcohol-related cancers^59^ because of limited availability of summary data. However, our gene-set enrichment analysis showed a significant enrichment for genes related to abdominal cancers.

Strengths of our study include its size, detailed attention to the alcohol phenotype, dense coverage of the genome through imputation, incorporation of brain and other imaging data to explore potential mechanisms and confirmatory *Drosophila* functional genetic studies. Over 80% of the data came from UKB, which combines high-quality phenotypic data and imputed genome-wide genetic data with strict attention to quality control^60^. We adopted a stringent approach to claim novel variants involving a conservative *P*-value threshold, internal replication in UKB and consistent direction of effect with the other studies, to minimize the reporting of false positive signals.

However, since alcohol intake is socio-culturally as well as genetically determined, it is influenced by other lifestyle and environmental factors which may modify or dilute the genetic signal. A key limitation is that assessment of alcohol intake relies on self-report, which is prone to errors and biases including recall bias and systematic under-reporting by heavy drinkers^61,62^. Furthermore, questionnaires on alcohol intake covered a short duration (e.g. day or week) at a single period, which may not be representative of broader drinking patterns of cohort participants. We harmonized data across cohorts by converting alcohol intake into a common metric of g/d, with imputation as necessary in UKB for participants reporting consumption of small amounts of alcohol. Taking this approach, we were able to detect strong genetic associations with alcohol intake that explained 7% of the variance in alcohol in an independent cohort, while our GRS analysis indicates that individuals in the lower fifth of the GRS distribution were consuming daily approximately one third of a standard drink (2.6 g/d alcohol) less compared with those in the upper fifth.

In summary, in this large study of genetic associations with alcohol consumption, we identified common variants in 46 novel loci with several of the genes expressed in the brain as well as other tissues. Our findings suggest that there may be common genetic mechanisms underpinning regulation of alcohol intake and development of a number of neuropsychiatric disorders including schizophrenia. This may form the basis for greater understanding of observed associations between excessive alcohol consumption and schizophrenia^63^.

## URLs

GTEx: www.gtexportal.org

UKBEC: http://www.braineac.org/

WebGetstalt: http://www.webgestalt.org

IPA: www.qiagen.com/ingenuity

PhenoScanner: http://www.phenoscanner.medschl.cam.ac.uk (Phenoscanner integrates results from the GWAS catalogue: https://www.ebi.ac.uk/gwas/ and GRASP: https://grasp.nhlbi.nih.gov/)

## Acknowledgements

H.G. was funded by the NIHR Imperial College Health Care NHS Trust and Imperial College London Biomedical Research Centre. I.K. was supported by the EU PhenoMeNal project (Horizon 2020, 654241) and the UK Dementia Research Institute, which is supported by the MRC, the Alzheimer‘s Society and Alzheimer’s Research UK. S.Thériault was supported by the Canadian Institutes of Health Research and Université Laval (Quebec City, Canada). L.R. was supported by Forschungs- und Förder-Stiftung INOVA, Vaduz, Liechtenstein. D.C. holds a McMaster University Department of Medicine Mid-Career Research Award. M.B. is supported by NIH grant R01-DK062370. P.v.d.H. was supported by ICIN-NHI and Marie Sklodowska-Curie GF (call: H2020-MSCA-IF-2014, Project ID: 661395). C.H. was supported by a core MRC grant to the MRCHGU QTL in Health and Disease research programme. N.V. was supported by Marie Sklodowska-Curie GF grant (661395) and ICIN-NHI. P.E. acknowledges support from the NIHR Biomedical Research Centre at Imperial College Healthcare NHS Trust and Imperial College London, the NIHR Health Protection Research Unit in Health Impact of Environmental Hazards (HPRU-2012-10141), and the Medical Research Council (MRC) and Public Health England (PHE) Centre for Environment and Health (MR/L01341X/1). P.E. is a UK Dementia Research Institute (DRI) professor, UK DRI at Imperial College London, funded by the MRC, Alzheimer’s Society and Alzheimer’s Research UK. This work received support from the following sources: the European Union-funded FP6 Integrated Project IMAGEN (Reinforcement-related behaviour in normal brain function and psychopathology) (LSHM-CT- 2007-037286), the Horizon 2020 funded ERC Advanced Grant ‘STRATIFY’ (Brain network based stratification of reinforcement-related disorders) (695313), ERANID (Understanding the Interplay between Cultural, Biological and Subjective Factors in Drug Use Pathways) (PR-ST-0416-10004), BRIDGET (JPND: BRain Imaging, cognition Dementia and next generation GEnomics) (MR/N027558/1), the FP7 projects IMAGEMEND(602450; IMAging GEnetics for MENtal Disorders) and MATRICS (603016), the Innovative Medicine Initiative Project EU-AIMS (115300-2), the Medical Research Council Grant ‘c-VEDA’ (Consortium on Vulnerability to Externalizing Disorders and Addictions) (MR/N000390/1), the Swedish Research Council FORMAS, the Medical Research Council, the National Institute for Health Research (NIHR) Biomedical Research Centre at South London and Maudsley NHS Foundation Trust and King’s College London, the Bundesministeriumfür Bildung und Forschung (BMBF grants 01GS08152; 01EV0711; eMED SysAlc01ZX1311A; Forschungsnetz AERIAL 01EE1406A, 01EE1406B), the Deutsche Forschungsgemeinschaft (DFG grants SM 80/7-2, SFB 940/2), the Medical Research Foundation and Medical research council (grant MR/R00465X/1), the Human Brain Project (HBP SGA 2). Further support was provided by grants from: ANR (project AF12-NEUR0008-01 - WM2NA, and ANR-12-SAMA-0004), the Fondation de France, the Fondation pour la Recherche Médicale, the Mission Interministérielle de Lutte-contre-les-Drogues-et-les-Conduites-Addictives (MILDECA), the Assistance-Publique-Hôpitaux-de-Paris and INSERM (interface grant), Paris Sud University IDEX 2012; the National Institutes of Health, Science Foundation Ireland (16/ERCD/3797), U.S.A. (Axon, Testosterone and Mental Health during Adolescence; RO1 MH085772-01A1), and by NIH Consortium grant U54 EB020403, supported by a cross-NIH alliance that funds Big Data to Knowledge Centres of Excellence.

## Author contributions

### Central analysis

E.E., H.G., C.C., G.N., P.B., A.R.B., R.P., H.Suzuki, F.K., A.M.Y., I.K., J.E., N.D., D.L., I.T., J.D.B., P.M.M., A.R., S.D., G.S., P.E.

### Writing of the manuscript

E.E., H.G., C.C., G.N., P.B., A.R.B., R.P., H.Suzuki, F.K., A.M.Y., I.K., D.L., I.T., J.D.B., P.M.M., A.R., S.D., G.S., P.E.

### Association of MRI analysis

C.C., H.Suzuki, A.M.Y., A.I.B., J.D.B., P.M.M., G.S.

### Alcgen and Charge+ contributor

(ARIC): A.C.M., M.R.B., B.Y., D.E.A., (CHS): B.M.P., R.N.L., T.M.B., J.A.B., (FHS): D.L., C.L., (GAPP/Swiss-AF/Beat-AF): S.Thériault, S.A., D.C., L.R., M.Kühne, (GENOA): S.L.R.K., J.A.S., W.A., S.M.R., (GRAPHIC): N.J.S., C.P.N., P.S.B., (GS): A.M.M., T-K.C., C.H., D.P., (HBCS): L.J., S.Tuominen, M.M.P., J.G.E., (HRS): D.R.W., S.L.R.K., J.D.F., W.Z., J.A.S., (MESA): X.G., J.Y., A.W., J.I.R., (METSIM): M.L., A.S., J.Vangipurapu, J.K., (FUSION): M.B., K.L.M., L.J.S., A.U.J., (NESDA): B.W.J.H.P., Y.M., (NFBC): M-R.J., J.Veijola, M.Männikkö, J.A., (ORCADES): H.C., P.K.J, (VIKING): J.F.W., K.A.K., (Croatia-VIS): I.R., O.P., (Croatia-KORCULA): C.H., (PREVEND): N.V., P.v.d.H, (OZALC): M.G.N., J.B.W., P.A.L., A.C.H., (SHIP): A.T., H.J.G., S.E.B., G.H., (TRAILS-pop): A.J.O, I.M.N., (TRAILS-CC): C.A.H., H.Snieder, (TwinsUK): T.D.S, M.Mangino, (YFS): L-P.L., M.Kähönen, O.T.R., T.L.

***All authors critically reviewed and approved the final version of the manuscript***

## Online Methods

### UK Biobank data

We conducted a Genome Wide Association Study (GWAS) analysis among 458,577 UKB participants of European descent, identified from a combination of self-reported and genetic data. The details of the selection of the participants has been described elsewhere^14^. These comprise 408,951 individuals from UKB genotyped at 825,927 variants with a custom Affymetrix UK Biobank Axiom Array chip and 49,626 individuals genotyped at 807,411 variants with a custom Affymetrix UK BiLEVE Axiom Array chip from the UK BiLEVE study, which is a subset of UKB. For our analyses, we used SNPs imputed centrally by UKB using the Haplotype Reference Consortium (HRC) panel.

#### Alcohol intake

We calculated the alcohol intake as grams of alcohol per day (g/d) based on self-reported alcohol drinking from the touch-screen questionnaire. The quantity of each type of drink (red wine, white wine, beer/cider, fortified wine, spirits) was multiplied by its standard drink size and reference alcohol content. Drink-specific intake during the reported drinking period (a week for frequent drinkers defined as: daily or almost daily/once or twice a week/three or four times a week; or a month for occasional drinkers defined as: one to three times a month/special occasions only) was summed up and converted to g/d alcohol intake for all participants with complete response to the quantitative drinking questions. The alcohol intake for participants with incomplete response was imputed by bootstrap resampling from the complete responses, stratified by drinking frequency (occasional or frequent) and sex.

Participants were defined as life-time non-drinkers if they reported ‘never‘ on the question on alcohol drinking frequency (UKB field 1558) and ‘no’ for the question on former drinker (UKB field 3731); they were excluded from further analysis. Participants with daily alcohol consumption > 500 grams we considered outliers and they were dropped from the analyses. We also excluded participants with missing covariates, leaving data on 404,732 individuals. We log_10_ transformed g/d alcohol and sex-specific residuals were derived from the regression of log_10_ transformed g/d alcohol on age, age^2^, genotyping chip and weight.

### UKB genetic analysis

We performed linear mixed modeling using BOLT-LMM software^64^, under an additive genetic model, for associations of measured and imputed SNPs with alcohol consumption (sex-specific residuals of the log_10_ transformed g/d variable). Model building was based on SNPs with MAF > 5%, call rate > 98.5% and HWE *P* > 1 × 10^-6^. SNPs were imputed using the HRC panel with imputation quality INFO score > 0.1. We estimated the LD score regression (LDSR) intercept to access the degree of genomic inflation beyond polygenicity as well as the lambda inflation factor λ_GC_^65^.

### The Alcohol Genome-Wide Consortium (AlcGen) and the Cohorts for Heart and Aging Research in Genomic Epidemiology Plus (CHARGE+) consortia

We analyzed available GWAS data from 25 independent studies (N=76,111) from the AlcGen and the CHARGE+ consortia. All study participants were of reported European ancestry and data were imputed to either the 1000 Genome Project or the HRC panel. Alcohol intake in g/d was computed and the log_10_ transformed residuals were analyzed as described above. Study names, cohort information and general study methods are included in Supplementary Table 2 and 3.

All studies were centrally quality-controlled using easyQC^66^. Finally, we analyzed data on ~7.1 M SNPs at MAF >1% and imputation quality score (Impute [Info score] or Mach [r^2^]) > 0.3. Genomic control (GC) was applied at study level. We synthesized the available GWAS using a fixed effects inverse variance weighted meta-analysis and summary estimates were derived for AlcGen and CHARGE+.

### One-stage meta-analysis

We performed a one-stage meta-analysis applying a fixed-effects inverse variance weighted meta-analysis using METAL^67^ to obtain summary results from the UKB and and the AlcGen plus CHARGE+ GWAS, for up to N=480,842 participants and ~7.1 M SNPs with MAF ≥ 1% for variants present in both the UKB data and AlcGen and CHARGE+ meta-analysis. The LDSR intercept (standard error), in the discovery meta-analysis was 1.05 and no further correction was applied.

### Previously reported (known) SNPs

We looked up in the GWAS catalog (http://www.ebi.ac.uk/gwas/) and identified 17 SNPs that associated with alcohol consumption at genome-wide significance level (*P* < 5 × 10^-8^). We enhanced the list by reference to a recent GWAS by Clarke et al^6^ that was not covered by the GWAS catalog at the time of the analysis, reporting 14 additional rare and common novel SNPs. Together with a SNP in RASGRF2 shown to be associated with alcohol-induced reinforcement^68^, we found 31 previously reported alcohol consumption related SNPs.

### Novel loci

According to locus definition of i) SNPs within ±500kb distance of each other; ii) SNPs in linkage disequilibrium LD (r^2^ > 0.1) calculated with PLINK, we augmented the list of known SNPs to all SNPs present within our data, not contained within the previously published loci. We further excluded SNPs in the HLA region (chromosome 6, 25-34Mb) due to its complex LD structure. We performed LD clumping in PLINK on 4,515 unknown SNPs with *P* < 1 ×10^-8^ using an r^2^ > 0.1 and distance threshold of 500kb. We further grouped the lead SNPs within 500kb from each other into the same loci and selected the SNP with smallest *P*-value from the locus as sentinel SNP. To report a SNP as novel signal of association with alcohol consumption:

i. the sentinel SNP has *P* < 5 × 10^-9^ in the one-stage meta-analysis;
ii. the sentinel SNP is strongly associated (*P* < 5 × 10^-7^) in the UKB GWAS alone;
iii. the sentinel SNP has concordant direction of effect between UKB and AlcGen and CHARGE+ datasets;
iv. The sentinel SNP is not located within any of the previously reported loci

We selected the above criteria i) to iii) to minimize false positive findings including use of a conservative one-stage *P*-value threshold that is an order of magnitude more stringent than a genome-wide significance *P*-value. (The threshold of *P* < 5 × 10^-9^ has been proposed e.g. for whole-genome sequencing-based studies.) This approach led us to the identification of 46 sentinel SNPs in total.

### Conditional analysis

We conducted locus-specific conditional analysis using the GCTA (Genome-wide Complex Trait Analysis) software (http://cnsgenomics.com/software/gcta). For each of the 46 novel sentinel SNPs, we obtained conditional analysis results for the SNPs with MAF>1% and within 500kb from the sentinel SNP after conditioning on the sentinel SNP. The meta-analysis results of the GWAS in UKB, AlcGen and CHARGE+ were used as input summary statistics and the individual-level genetic data from UKB were used as the reference sample. Results for a SNP were considered conditionally significant if the difference between the conditional *P*-value and the original *P*-value is greater than 1.5-fold (-log_10_*P*/-log_10_(*P*_conditional) >1.5 and the conditional *P*-value is smaller than 5 × 10^-8^.

### Gene expression analyses

To analyze the impact of genetic variants on expression of neighboring genes and identify expression quantitative trait loci (*cis*-eQTLs; i.e., SNPs associated with differences in local gene expression), we used two publicly available databases, the Genotype-Tissue Expression (GTEx) database^69^ and the UK Brain Expression Consortium (UKBEC) dataset^70^. We searched these databases for significant variant-transcripts pairs for genes within 1Mb of each input SNP.

With the GTEx database, we tested for *cis*-eQTL effects in 48 tissues from 620 donors. The data described herein were obtained from the GTEx Portal, Release: V7 and used FastQTL^71^, to map SNPs to gene-level expression data and calculate q-values based on beta distribution-adjusted empirical *P*-values^72^. A false discovery rate (FDR) threshold of ≤0.05 was applied to identify genes with a significant eQTL. The effect size, defined as the slope of the linear regression, was computed in a normalized space (normalized effect size (NES)), where magnitude has no direct biological interpretation. Here, NES reflects the effects of our GWAS A1 alleles (that are not necessarily the alternative alleles relative to the reference alleles, as reported in the GTEx database). Supplementary Table 12 lists transcripts-SNPs associations with significant eQTL effects.

With the UKBEC dataset that comprises 134 brains (http://www.braineac.org/), we searched for *cis*-eQTLs in 10 brain regions, including the cerebellar cortex (CRBL), frontal cortex (FCTX), hippocampus (HIPP), medulla (specifically inferior olivary nucleus, MEDU), occipital cortex (specifically primary visual cortex, OCTX), putamen (PUTM), substantia nigra (SNIG), thalamus (THAL), temporal cortex (TCTX) and intralobular white matter (WHMT), as well as across all brain tissues (aveALL). MatrixEQTL^73^ generated *P*-values for each expression profile (either exon-level or gene-level) against the respective SNP were obtained for the 10 different tissues and overall (aveALL). **Supplementary Table 13** lists transcripts-SNPs associations with a eQTL *P*-value < 0.0045 in at least one brain tissue. Subsequent data analysis was performed in R (http://www.R-project.org/).

We carried out over-representation enrichment analysis using the list of 146 GTEx eQTL genes. Ingenuity pathway analysis (IPA^®^, QIAGEN Inc.) was performed on this list using ontology annotations from all available databases except those derived from low-confidence computational predictions.

### Magnetic Resonance Imaging Data

We used the most recent release of magnetic resonance imaging (MRI) data on brain, heart and liver for UKB participants to investigate genetic associations with the 46 novel SNPs for alcohol consumption.

### Brain imaging

#### Brain MRI acquisition and pre-processing

We used the T1 data from UKB to elucidate volumetric brain structures, including the cortical and the sub-cortical areas. The T1 data were acquired and pre-processed centrally by UKB. The brain regions were defined by combining the Harvard-Oxford cortical and subcortical atlases^74^ (https://fsl.fmrib.ox.ac.uk/fsl/fslwiki/Atlases) and the Diedrichsen cerebellar atlas^75^ (http://www.diedrichsenlab.org/imaging/propatlas.htm). FAST (FMRIB’s Automated Segmentation Tool)^76^ was then used to estimate the grey matter partial volume within each brain region. Subcortical region volumes were also modelled by using FIRST (FMRIB’s Integrated Registration and Segmentation Tool). More details about the MRI scanning protocol and pre-processing has been provided in UKB documentation (https://biobank.ctsu.ox.ac.uk/crystal/docs/brain_mri.pdf).

#### Association Analyses

We performed association analyses on N = 9,705 individuals between all novel SNPs and the grey matter volume of brain regions using Pearson correlation, adjusting for age, age^2^, sex, age × sex, age^2^ × sex, and head size. All, brain volume features, log transformed alcohol intake data (g/d), and the confounders were firstly transformed by using a rank-based inverse Gaussian transformation. Significance levels were set at *P* < 0.05 adjusted using the false-discovery rate method for multiple comparisons.

#### Mediation analysis

To assess if the effect of a SNP on alcohol consumption is mediated through a brain region, we performed a single-level mediation analysis based on a standard three-variable path model (SNP-brain region-alcohol consumption) with corrected and accelerated percentile bootstrapping 10,000 times to calculate the significance of the mediation effect. We considered as mediator variable the grey matter volume of brain regions that had a significant association on alcohol consumption. We calculated the significance of path a, path b and a*b mediation (SNP-brain region-alcohol consumption) using a multilevel mediation and moderation (M3) toolbox^77,78^

### Cardiac Imaging

#### Cardiac MRI acquisition and pre-processing

Details of the cardiac image acquisition in UKB are reported previously^79^. Cardiac MRI was acquired using a clinical wide bore 1.5T scanner (MAGNETOM Aera, Syngo Platform VD13A, Siemens Healthcare, Erlangen, Germany) with 48 receiver channels, a 45 mT/m and 200 T/m/s gradient system, an 18-channel anterior body surface coil used in combination with 12 elements of an integrated 32 element spine coil and electrocardiogram gating for cardiac synchronization. A two-dimensional short-axis cardiac MRI was obtained using a balanced steady state free precession to cover the entire left and right ventricle (echo time, 1.10msec; repetition time, 2.6msec; flip angle, 80°; slice thickness, 8mm with 2mm gap; typical field of view, 380×252mm; matrix size, 208×187, acquisition of 1 slice per breath-hold).

The cardiac images were segmented to provide left ventricular mass (LVM), left end-diastolic (LVEDV), left end-systolic volume (LVESV), and right end-diastolic (RVEDV) and right end-systolic volume (RVESV) using a fully convolutional network as described previously^80^. Left (LVEF) and right ventricular ejection fraction (RVEF) were derived from (LVEDV–LVESV)/LVEDV×100 and (RVEDV– RVESV)/RVEDV×100, respectively.

#### Association Analyses

To test associations between cardiac MRI measures and alcohol consumption-related SNPs, we carried out a regression of LVM, LVEDV, LVEF, RVEDV, and RVEF onto each of the 46 SNPs adjusting for age, sex, height, weight, hypertension, diabetes, and smoking history. Significance levels were set at *P* < 0.05 adjusted using the false-discovery rate method for multiple comparisons.

### Liver Imaging

#### Liver MRI acquisition and pre-processing

Details of the liver image acquisition protocol have been reported previously^81^. Briefly, all participants were scanned in a Siemens MAGNETOM Aera 1.5-T MRI scanner (Siemens Healthineers, Erlangen, Germany) using a 6-minute dual-echo Dixon Vibe protocol, providing a water and fat separated volumetric data set for fat and muscle covering neck to knees. For liver proton density fat fraction (PDFF) quantification, an additional single multi-echo gradient slice was acquired over the liver. Liver images were analysed by computing specific ROI for water, fat and T2* by magnitude-based chemical shift technique with a 6-peak lipid model, correcting for T1 and T2*

#### Association Analyses

We performed association analyses between 46 alcohol consumption-related SNPs and liver PDFF (%), from 8,372 samples, using a linear regression model adjusting for age, age^2^, sex, T2D, BMI, genotyping chip and first three PCs. Liver PDDF was firstly transformed by using a rank-based inverse transformation. Significance levels were set at *P* < 0.05 adjusted using the false-discovery rate method for multiple comparisons.

### *Drosophila* experiments

Flies were kept on standard cornmeal/molasses fly food in a 12:12hr light:dark cycle at 25°C. Transgenc flies were obtained from the Bloomington *Drosophila* Stock Center: *UAS-hZip8* BL#66125, *UAS-dZIP71B-TRiP-RNAi^HMC04064^* BL#55376, *dZip71B^MI13940^* BL#59234, and *dZip71B^MB11703^* BL#29928. For behavioral experiments, crosses were set up such that experimental and control flies were sibling progeny from a cross, and both were therefore in the same hybrid genetic background (*w Berlin / unknown*). Flies aged 1-5 days of adult age were collected, exposed to 100/50 (flowrates) ethanol/air vapor in the Booze-o-Mat 2 days later, and their loss of righting determined by slight tapping, as described^82^. For tolerance, flies were put back onto regular food after a 30-min initial exposure, and were then re-exposed to the same vapor 4 hours later. Note that tolerance is not connected to initial sensitivity, and flies naively sensitive to ethanol-induced sedation can have no, or a reduced tolerance phenotype. Flies overexpressing *hZip8* (and their sibling controls) were placed at 28°C for two days to increase the expression levels of the transgene, as we did not detect a phenotype when they were kept at 25°C (data not shown). Data from experimental and control flies were compared by Student’s t-tests.

### Effects on other traits and diseases

We queried SNPs against GWAS results included in PhenoScanner, to investigate cross-trait effects, extracting all association results with genome-wide significance at *P* < 5 × 10^-8^ for all SNPs in high LD (r^2^ ≥ 0.8) with the 46 sentinel novel SNPs, to highlight the loci with strongest evidence of association with other traits. At the gene level, overrepresentation enrichment analysis (ORA) with WebGestalt^41^ on the nearest genes to all alcohol intake loci was carried out.

The genetic correlations between alcohol consumption and 235 other traits and diseases were obtained in the online software LD Hub. LD hub is a centralized database of summary-level GWAS results and a web interface for LD score regression analysis To estimate the potential causal effect of alcohol consumption-related variants on schizophrenia, we performed a Mendelian randomization analysis utilizing publicly available GWAS data on schizophrenia and the Mendelian randomization package in R. The effect was estimated using the inverse-variance weighted (IVM) method. Pleiotropy was tested by applying the MR-Egger regression method and heterogeneity statistics were obtained. In presence of heterogeneity the random effects inverse-variance method was applied^83^.

### Genetic risk scores and percentage of variance explained

We calculated an unbiased weighted genetic risk score in 14,004 unrelated participants in Airwave, an independent cohort with high quality HRC imputed genetic data^33^. We used as weights the beta coefficients of the meta-analysis. We assessed the association of the GRS with alcohol intake and calculated the alcohol consumption levels for individuals in the top vs the bottom quintiles of the distribution. To calculate the percent of variance of alcohol consumption explained by genetic variants, we generated the residuals from a regression of alcohol consumption in Airwave. We then fit a second linear model for the trait residuals with all novel and known variants plus the top 10 principal components, and estimated the percentage variance of the dependent variable explained by the variants.

